# The role of fecundity dynamics and time of injury for female reproductive fitness

**DOI:** 10.1101/2025.11.12.687561

**Authors:** Piotr Michalak, Jean-Baptiste Ferdy, David Duneau

**Affiliations:** Université de Toulouse, Toulouse INP, CNRS, IRD, CRBE, Toulouse, France; The Queen’s Medical Research Institute, Centre for Cardiovascular Science, Edinburgh University, Edinburgh, UK; Center for Ecology, Evolution and Environmental Changes (cE3c) & Global Change and Sustainability Institute (CHANGE), Faculty of Sciences, University of Lisbon (FCUL), 1749-016 Lisbon, Portugal

**Keywords:** oviposition, injury, fecundity, dynamics, *Drosophila melanogaster*

## Abstract

Animals have a wide range of reproductive strategies and change how they produce offspring as a function of their age, environment, risk of predation or starvation, and as a result of trade-offs with other life history traits. In turn, these temporal patterns of reproduction, or fecundity dynamics, will determine how vulnerable the animal is to many of the same factors that shaped the dynamics in the first place. In this study, we tested how individual fecundity dynamics influence vulnerability of female fecundity to the deleterious environmental factors. We injured *Drosophila melanogaster* females at various moments during periods before and after mating. Injuries are prevalent in nature and provide a punctuated stimulus, making them particularly well suited to study their connection with the dynamics of reproduction. Our aim was to determine periods of vulnerability and impact of wounds on female fitness resulting from the change. The observed fecundity dynamics exhibited a high early peak in offspring production, suggesting that early after mating females might be particularly vulnerable to environmental conditions. However, females produced the same number of offspring, regardless of the time of injury, suggesting that fitness of female *Drosophila* is robust to wounds inflicted under laboratory conditions.

## 1 Introduction

Animals and plants have adapted to various environments using a wide range of reproductive strategies, which are the result of trade-offs between fecundity and other life history traits [1–4]. These strategies include fast or slow reproduction; reproducing continuously or seasonally; maintaining steady pace of offspring production or producing offspring in bursts [2, 3]. However, regardless of the reproductive strategy, reproducing animals face common challenges, including infections, ectoparasitism, injury due to competition with conspecifics, and encounters with predators [5–8]. The vulnerability of females’ fertility to those conditions, many of which might necessitate redirecting essential resources away from reproduction toward other critical functions, such as defence, escape behaviours, wound healing, and immune responses to infection, has a direct implication for evolutionary fitness. This vulnerability further varies with female’s current reproductive potential: older individuals who have finished reproducing incur no reproductive cost regardless of the challenge, while females at the peak of their fecundity face disproportionately high cost when diverting the resources from reproduction. Female current reproductive state is therefore a major factor which determines her fitness vulnerability.

In most species, as fecundity generally declines with age, the most vulnerable stage in a female’s lifespan should be the onset of her reproductive life [3]. However, in sperm-storing organisms, such as many insects and birds, this pattern of peak vulnerability also occurs at shorter timescales within the reproductive period. In *Drosophila*, just after mating, females have a peak of fecundity that declines as available sperm depletes, until the female runs out of sperm or remates [9, 10]. Consequently, a challenge occurring immediately after mating will divert resources when they are most needed for reproduction, likely leading to the largest fitness cost. However, these changes in fecundity over time, or fecundity dynamics, are often overlooked when total fitness is measured.

*Drosophila melanogaster* provides an ideal system for testing how vulnerability of reproduction varies with fecundity dynamics. The fecundity dynamics of this species are well understood, particularly across an individual’s lifetime [11–14, among others], and females can store sperm for over a week, providing a window into the changing fecundity after mating. Importantly, females of this species are naturally exposed to challenges that can divert resources from reproduction: they frequently suffer injuries from ectoparasite [8], which can then lead to infection and initiate a costly immune response [15, 16], though even a sterile injury alone can decrease offspring production [17]. This combination of a well-characterized fecundity pattern, natural reproductive challenges, and extended sperm storage makes *D. melanogaster* particularly suitable for examining how the timing of challenges relative to peak fecundity, affects reproductive costs.

In this study, we studied how fecundity dynamics influence the fitness costs associated with injury in *Drosophila melanogaster*. Because sterile injury is a punctuated challenge, it is well suited to study its impact on the dynamics of reproduction. Furthermore, experimental injuries are known to be costly as they notably reduce the chances to survive subsequent infections [18, 19]. We tracked single-mated, individually housed *D. melanogaster* females and followed their daily offspring production. Using only single mating provides access to the full post-mating dynamics as related to sperm depletion, and gives a complete picture of the fitness costs of injury. It also removes potential effect of injury on remating propensity. We conducted two experiments to test the hypothesis that reproductive vulnerability varies with fecundity. First, we wounded females with a glass needle before and after mating to simulate abdominal injuries they might frequently sustain in nature. We predicted that post-mating females would show greater costs due to their higher fecundity. Second, we wounded females at different times after mating. We hypothesized that the costs of injury would be highest immediately after mating (fecundity peak) and would decrease as fecundity declines over time.

## 2 Methods

### 2.1 Effects of wounding pre- and post-mating on offspring production

*Drosophila melanogaster* individuals (DGRP genotype RAL21) were reared on commonly used food medium (corn flour 74 g/l, yeast 28 g/l, sugar 40 g/l, agar 8 g/l, Moldex (20% OH) g/l, propionic acid 3 g/l) in a 12hr light/dark cycle at 25°C. To prevent overcrowding during development, parents were placed in 12 vials with approximately eight females and four males per vial for two days.

We used a three-treatment design to test the effects of injury timing on reproductive output: (1) injury before mating, (2) injury after mating, and (3) uninjured controls.

#### Day 0

85 virgin females (4 days old, n=39; 6 days old, n=46) were randomly assigned to treatments at the start of the experiment. Twenty eight individuals assigned to pre-mating injury group were wounded in the abdomen using a sterile glass needle under CO_2_ anaesthesia. All flies were then housed individually in rearing fly vials with randomized rack positions to prevent block effects.

#### Day 2

All females were placed, in the morning, with males for mating. Females were flipped into new vials containing one male and allowed to mate for eight hours. Mating duration was recorded, and males were removed after copulation. Thirteen females that did not mate within this time-frame were left with males overnight, and males were removed the following morning. If those females produced offspring, the dynamics were included in the dataset as the patterns of sperm depletion are not expected to be different.

#### Day 4

From the remaining unmated females, 28 previously selected were wounded (post-mating injury group). Twenty nine females served as uninjured controls.

Following mating, females were transferred daily to fresh vials for 14 consecutive days. To improve larval survival at low densities, a small hole was made in each food medium to facilitate larval access. After approximately 14 days post-laying (with additional days monitored to ensure complete emergence), all adult offspring were counted to determine daily fecundity.

The final dataset included 59 females after removing non-virgin individuals, escapees, and females that produced no offspring despite observed mating (n=13). Sample sizes by treatment were: control (n=22), pre-mating injury (n=17), and post-mating injury (n=20). Females that died of natural causes during the experiment were included in analyses up to their death date, since they were considered to have finished their reproduction. Some vials across treatments showed fungal contamination, but this was not systematically recorded.

### 2.2 Effects of wounding at different times post-mating

*D. melanogaster* were reared as described in the previous experiment. We used two genotypes to introduce some controlled genetic variability: DGRP RAL21 (parents were placed in four vials; three vials with eight females and four males, one of six females and three males) and DGRP RAL783 (six vials of eight females and four males).

#### Day -4 (within 8 hours post-eclosion)

Virgin flies were separated by sex under CO_2_ anaesthesia and moved into new vials.

#### Day -1

Females were placed under anaesthesia and housed individually in vials.

#### Day 0 (4 days post-eclosion)

Females were placed with males and allowed to mate for eight hours. Mating duration was recorded, and after copulation, males were removed. A total of 95 females mated successfully (RAL21=42, RAL783=53).

#### Days 1-3 post-mating

Females were randomly assigned to four treatment groups: wounding on day 1 post-mating (n=23), day 2 post-mating (n=25), day 3 post-mating (n=23), or unwounded controls (n=24). Wounding in the abdomen was performed using a sterile glass needle under CO_2_ anaesthesia.

Following mating, females were transferred each morning into fresh vials for 14 days. As in the previous experiment, a hole was made in the food medium to facilitate larval survival. Adult offspring were counted approximately 14 days after laying (with additional monitoring to ensure complete emergence).

Nineteen flies were excluded, either because they escaped or they died due to mishandling, in addition to four flies that did not reproduce. Flies that naturally died during the experiments were considered to have finished their reproduction and were included in the dataset. The final dataset consisted of 72 females (control, n=18; day 1, n=16; day 2, n=20; day 3, n=18) (see also Supplement).

Some vials developed some fungal contamination on the food during the experiment. An analysis showed that the appearance of fungus was associated with low number of offspring, probably because small number of larvae might not be sufficient to limit fungal growth. This is corroborated by the observation that the fungus had developed mainly later in the observation period, when fecundity was low. However, fungus was not a statistically significant predictor in the models used (see Supplement).

### 2.3 Statistical analyses

We used generalized linear models to analyse three response variable: total offspring production, peak fecundity and time it took to reach 50% of offspring production. As these variables are positive numbers that correspond to either number of offspring or specific day, we assumed a Poisson distribution and allowed for overdispersion by using a “quasipoisson” distribution. Our primary hypothesis was that the reproductive costs is affected by timing of injury relative to mating. We therefore focused our analysis on comparing the two injury treatments (before vs. after mating) to test whether pre-mating injury causes greater reproductive costs than post-mating injury, as predicted by our fecundity dynamics hypothesis. The model included wound timing, female age, and their interaction. The model reads as:

TOTAL REPRODUCTION or TIME TO 50% or PEAK REPRODUCTION *∼* WOUND_TIMING + AGE + WOUND_TIMING:AGE

For analysing the effect of injury on the duration of the copulation (measured in minutes), we used a GLM with a “gamma” distribution. This analysis compared flies wounded before mating, with uninjured flies at the time of mating (control and post-mating wounding groups).

MATING DURATION *∼* INJURED_STATUS + AGE + INJURED_STATUS:AGE

In the second experiment, we analysed the same variables as in the first experiment, but included additional predictors: mating duration, genotype, and fungus contamination status. Our hypothesis was that the injury cost would vary depending on timing post-mating, with earlier injuries causing greater costs. The model included all four treatment groups to test both injury effects and their temporal dynamics:

TOTAL REPRODUCTION or TIME TO 50% or PEAK REPRODUCTION *∼* WOUND_TIMING*GENOTYPE + GENOTYPE*MATING_DURATION + GENOTYPE*FUNGUS

A model including all possible interactions has also been tested (see Supplement).

We also quantified the recovery from the injury by contrasting fecundity one day post treatment to that in the following day. If injured flies are recovering, they should show positive changes through days, while control flies should show negative changes due to natural sperm depletion. We assumed that this difference has a normal distribution, and tested the effect of day of injury and genotype.

OFFSPRING DIFFERENCE *∼* (INJURED_STATUS + WOUND_DAY + GENOTYPE)^3^

where INJURED_STATUS contrasts injured vs. control flies, WOUND_DAY indicates timing of injury (day 1, 2, or 3 post-mating), and GENOTYPE accounts for line differences. Significance testing was based on the analysis of deviance in R “stats” package (for full table see Supplement). Data were analysed using R v.4.5.1 [20].

## 3 Results

### 3.1 Effects of wounding pre- and post-mating on female fecundity

The injury had no effect on copulation time (F_1,44_ = 3.95, p = 0.053, *≈* 3.15 minutes mean increase in duration of the copulation from 29.18 minutes duration in non-wounded flies). Copulation time did not differ between age at mating either (F_1,43_ = 1.51, p = 0.23, Fig. 1A).

**Figure 1.**
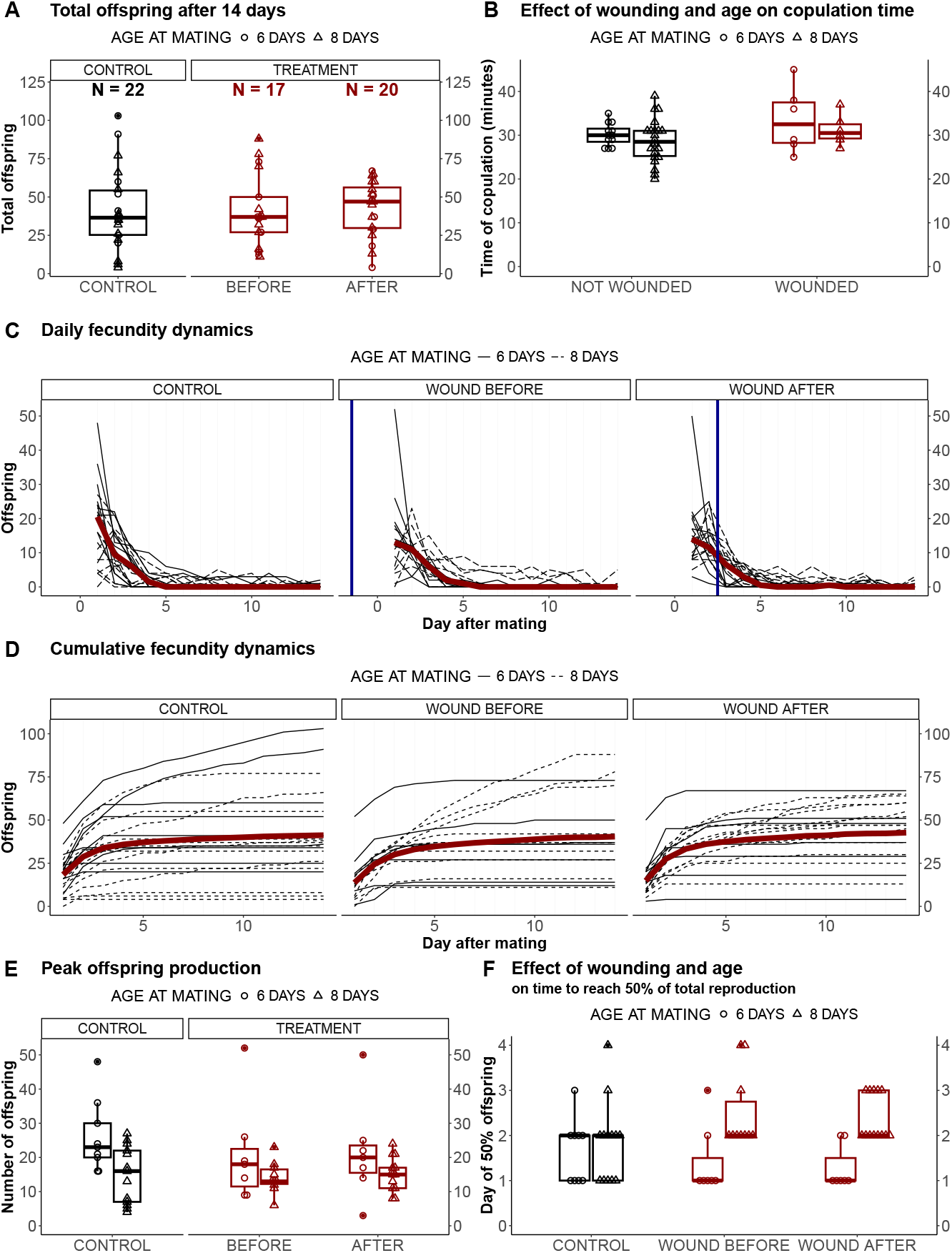
**A) Total fecundity after 14 days.**The treatment groups show a similar number of offspring. **B) The duration of copulation** The wounding before mating or age does not influence the duration of copulation. **C) Daily fecundity of *Drosophila* over a 14-days period**. Each black line represents a female that produced at least one offspring. The median number of offspring per day is shown in red. Period post treatment is marked by the blue vertical line. The dynamics exhibit a pronounced peak, followed by a decline and a period of low offspring production. **D) Cumulative fecundity over a 14-days period**. Most flies reproduce fast early on, then the number of offspring stabilizes or flies produce few offspring per day. The red line is the daily mean. **E) Maximal daily offspring production**. The peak offspring production accounts for an average of 50% of total offspring production. Age at mating impacts the height of the peak, with 8-day-old flies having lower peaks than 6-day-old. **F) Day at which 50% of offspring has been reached**. Injury does not change the speed of reproduction, but older flies reach the 50% of the reproduction later. A majority had produced 50% of their fitness within 2 days.

The total number of offspring produced during the experimental period varied from four to 103, demonstrating a considerable individual variation (mean= 41.49 *±* 22.58). We found no difference in this total offspring production between flies wounded before and flies wounded after mating (F_1,35_ = 0.11, p = 0.75, Fig. 1B) or between the two age groups (F_1,34_ = 1.62, p = 0.22). Since wounded flies that mated when 8 days old had lower peaks than flies mated at 6 days old but have the same total fecundity, it is possible that the older flies were able to compensate by producing offspring later in the experiment. Note that because we did not replicate the age treatment, age serves as a descriptive distinction rather than a statistically testable factor in our design.

The fecundity dynamics obtained in the experiments are characterized by an early peak in the daily offspring count, which occurred within 48 hours post mating (Fig. 1C). After the peak, daily offspring production quickly declined (Fig. 1C,D). The peak is an important feature of the dynamics, accounting on average for 50.1% *±* 21.5% of the total reproduction. There was no difference in the height of the peak between the treatments (F_1,35_ = 0.02, p = 0.88, Fig. 1E). However, the two age groups within treatments showed pronounced differences in the height of the peak (F_1,34_ = 4.65, p = 0.04) with older flies (8 days old at time of mating) having lower peaks (Fig. 1E). The two injury treatments also showed the same speed of reproduction, measured by the day a fly reached 50% of her total reproductive output (F_1,35_ = 0.07, p = 0.8, Fig. 1F). We observed that older wounded flies reached this 50% output later than younger wounded flies (F_1,34_ = 23.18, p = 0.00003, Fig. 1F).

### 3.2 The effects of injury at different times post-mating

The observed fecundity dynamics are similar to those described in the previous experiment (Fig 2C,D). They are characterized by a peak early on, followed by a rapid decline. The total offspring count after 14 days of laying was not different between the treatment groups (F_2,51_ = 0.76, p = 0.47, Fig 2C). We found a difference in the number of produced offspring between the two DGRP lines (F_1,50_ = 9.70, p = 0.003). However, there was an effect of interaction between duration of copulation and the DGRP line (F_1,45_ = 5.22, p = 0.03). We found no difference between the three treatments in the day they have reached 50% of the total reproduction (F_2,51_ = 1.24, p = 0.30, Fig 2D). Because several treatments happened post-mating and 50% might have been reached beforehand, we also considered the time flies reached 95% of the total reproduction to check the consistency. Here again, there was no difference between the treatments (F_2,51_ = 0.33, p = 0.72, Fig 2E).

**Figure 2.**
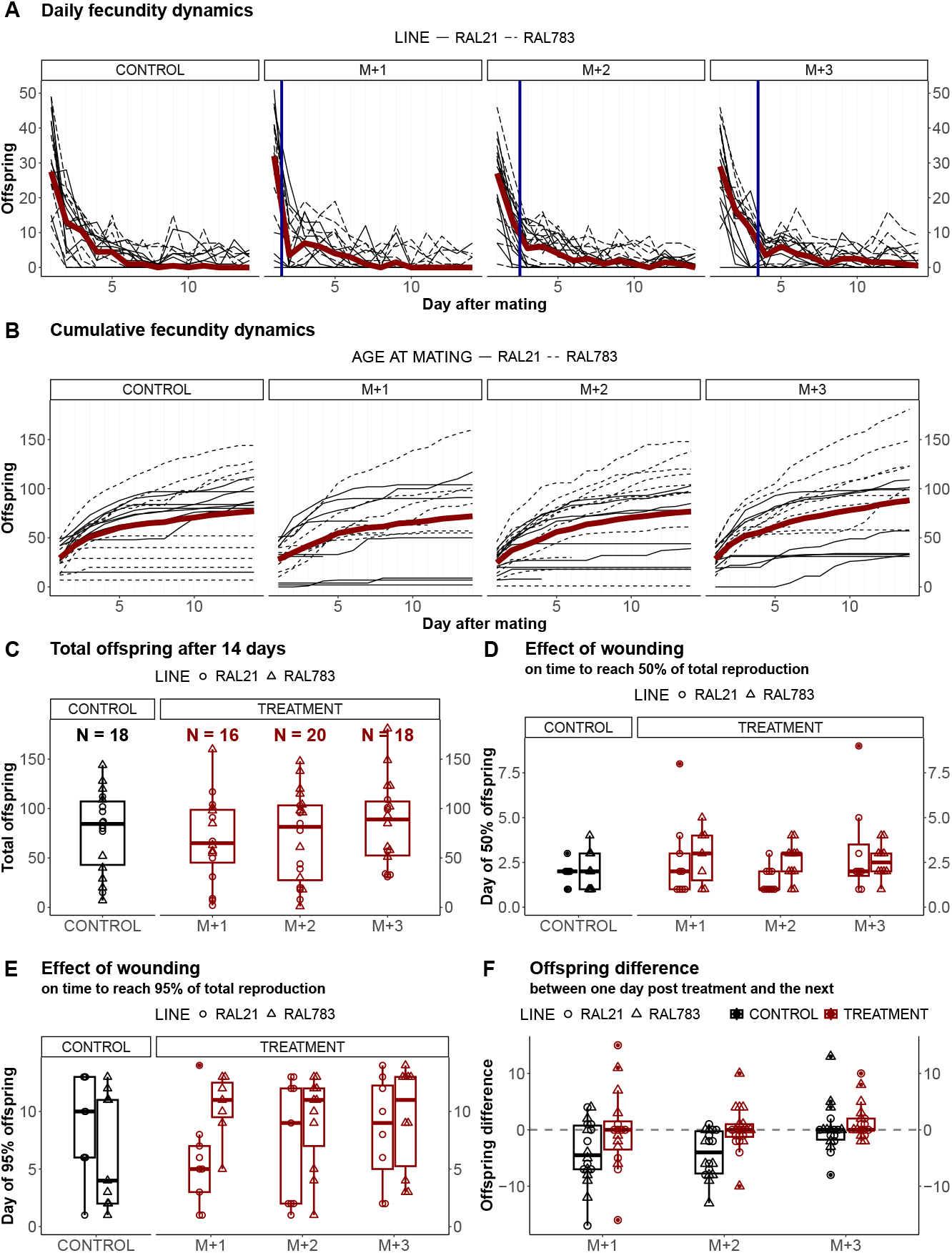
**A) Daily fecundity of *Drosophila***.Each line represents a female that produced at least one offspring. The median number of offspring per day is shown in red. The dynamics exhibit a pronounced peak, followed by a decline and a later period of low offspring production. On the day after treatment (blue vertical line), the reproduction appears to be lower than on the next day, suggesting a recovery after the injury. **B) Cumulative fecundity over a 14-days period**. The cumulative data show that most flies reproduce fast early on, then offspring production stabilizes. However, some flies continued to produce offspring even after 14 days. The red line represents the daily mean. **C) Total fecundity after 14 days**. The three treatments show similar number of offspring, suggesting that if injury influenced offspring production shortly after the wound had occurred, flies can still lay all the eggs they could from one mating. **D) Day at which 50% of offspring has been reached**. The wounded treatments do not differ in the speed of reproduction. **E) Day after mating at which 95% of offspring has been reached**. The treatments do not differ in the time to reach 95% of the total offspring production. Majority of the offspring that could be produced from one mating were laid within 10 days. **F) The difference between the control and wounded groups in the offspring difference between production one day after wound and the next**. The injury treatments differ in the number offspring produced between first and second day after treatment than corresponding days in the control, suggesting that the wound influenced offspring production.

Fig 2A suggests that daily offspring production drops the day immediately following the injury, but increases the day after, contrasting with the apparent steady decline in control flies. Analysis of day-today changes confirmed that treated flies were more likely to show this increase than control flies (F_1,105_ = 11.27, p = 0.001, Fig 2F). However, this pattern must be interpreted cautiously. First, this analysis was conducted post-hoc after observing the dynamics rather than being pre-planned. Second, control flies were not exposed to CO_2_ anaesthesia as it trade-offs with the total sample size we could handle in one experiment. Although, injured flies were anaesthetised for less than a minute, and it is not expected to have an impact on offspring production in our experimental conditions ([21]), we cannot exclude the possibility that the apparent recovery could reflect delayed egg-laying from anaesthesia rather than biological recovery from injury. Future experiments should test the hypothesis of a recovery and include anaesthetised-only control flies.

## 4 Discussion

By following individual females, we obtained individual dynamics of reproduction. As expected, these dynamics were characterized by a pronounced fecundity peak occurring on the first or second day after mating, followed by a rapid decline. This decline was likely attributable to the depletion of sperm in storage. Yet, despite low fecundity after five days, many flies continued to produce offspring until day 14, indicating that sperm were not fully depleted by the end of the experiment. However, contrary to our hypothesis, we found no effect of the time of injury around mating on total offspring production.

The lack of detectable decrease in offspring production due to injury is unexpected. Injury generally decreases total offspring production in *Drosophila* [17], and non-penetrative blunt force trauma has been showed to lead to egg retention and ovarian cell death [22, 23]. However, flies can mount an immune responses to heal sterile wounds, and such injury usually does not substantially decrease females’ survival [24]. Additionally, many trade-offs with fecundity arise only under specific conditions, such as calorie limitation [25]. It has also been argued that trade-offs with immunity in *Drosophila* are primarily from reduced energy availability due to decreased metabolism and food intake [26] and not due to energy reallocation to immune responses. Since we renewed the food medium daily, it is unlikely that females were limited by energy availability. As result, females may have had enough resources to both heal and maintain high offspring production, making treatment differences undetectable. Different conditions or more severe stressors might be needed to reveal trade-offs affected by the time of reproduction.

The early fecundity peak in *Drosophila* represents a large fraction of the total reproduction in the study. Since females enter a period of low fecundity around five days after mating, failing to account for the day of mating likely introduces considerable variation in some studies, as remating is unlikely to be synchronous in populations where females are co-housed with males. Even accounting for the time of mating time, as in our study, we observed considerable heterogeneity in the number of offspring produced. Some females never produced offspring despite mating [see also 13]. The resulting heterogeneity would be hidden in traditional group experiments where multiple females lay eggs together in a single vial [e.g. 11]. This supports the fact that studies should be cautious when reporting “eggs per fly” instead of “eggs per group of flies”, as an average number of egg laid by several co-housed flies is not biologically equivalent to that of a single fly. Moreover, studies that focus only on the reproductive output over a period of two or three days may miss critical effects that occur specifically during peak fecundity, potentially overlooking biologically meaningful responses to experimental treatments. However, while individual tracking provides clearer dynamics, it has drawbacks. *Drosophila* produces less offspring in the absence of adult conspecifics [27] and larval survival is suboptimal under low larval density. Thus, the cost-benefit analysis between individual and group egg laying experiments requires close considerations during experiment planning.

Although both fly groups were made of young flies, relatively older, wounded flies had lower fecundity peaks than wounded, younger flies. They were also slower to reach the 50% of their total offspring production. Despite these differences in pattern of reproduction, we found no difference in total offspring produced by the two age groups at the end of the experiment. Although we used only a single genotype and the groups differed in age by only two days, which is unlikely to cause pronounced effects, this pattern shows that early reduction in reproduction can be compensated later during the reproductive period. This compensation mechanism might be particularly relevant, given that some experiments measure only early fecundity [e.g. 28], potentially missing important biological responses and fitness consequences.

Our study presents a complex picture of reproductive vulnerability. On one hand, we confirmed that certain periods of fecundity dynamics, particularly the early peak, play a disproportionately important role in total fecundity of singly mated flies. However, we did not detect the expected differences in fitness costs based on the time of injury during these critical periods. Our underlying assumption was that injury costs would be immediate and irreversible, leading to a missed reproduction opportunity. This assumption aligns with evidence that females lose sperm viability or dump sperm when mounting immune responses [29]. However, it is possible that females shift their reproductive patterns and largely compensate for the early decrease over time. Given that wounding can decrease overall fecundity [17], how the time of this decrease changes the related fitness costs remains an open question in understanding life-history trade-offs.

## 5 Acknowledgements

We thank Christian Faucher, Nathalie Parthuisot and Aurore Gavenc for regularly helping with the experiments. Jennifer Perry and Mariana Wolfner for their valuable comments on the manuscript. P.M. was supported by PhD funding from Université Toulouse 3 – Paul Sabatier through the SEVAB doctoral school. D.D. was supported by a FCT fellowship (2023.08149.CEECIND).

## Authors’ contributions

P.M., D.D. and J.B.F. conceptualized the study; P.M. collected the data, analyzed the results and led the writing of the manuscript with assistance from D.D. and J.B.F.; All authors contributed critically to the drafts and gave their final approval for publication.

## Supplement

### 5.1 Effects of wounding pre- and post-mating on offspring production

#### 5.1.1 Flies that died or did not produce offspring - by treatment

| Treatment | N | Died (but produced offspring) | No offspring |
| --- | --- | --- | --- |
| Control | 25 | 0 | 3 |
| Wound after | 23 | 1 | 3 |
| Wound before | 24 | 1 | 7 |

Flies that did not produce offspring were not included in the analysis.

#### 5.1.2 Sample sizes used in the analyses - by age and treatment

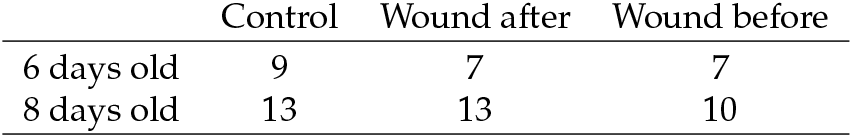

Note that the majority of the analyses is between treatments comparison.

#### 5.1.3 Full model results – ANOVA

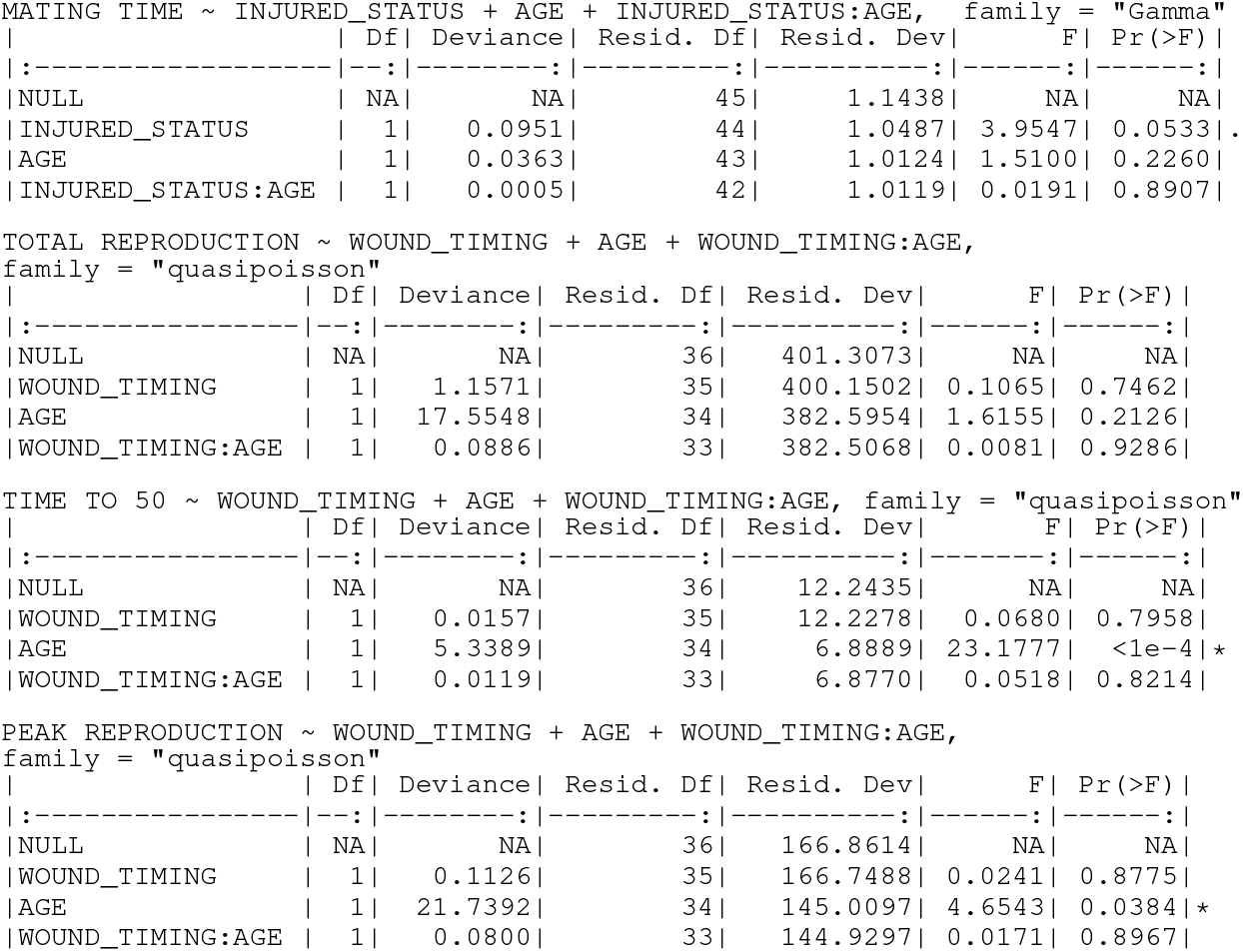

### 5.2 Effects of wounding at different times post-mating

#### 5.2.1 Flies that died or did not produce offspring - by treatment

Flies that did not produce offspring were not included in the analysis.

#### 5.2.2 Sample sizes used in the analyses - by line and treatment

Note that the majority of the analyses is between treatments comparison.

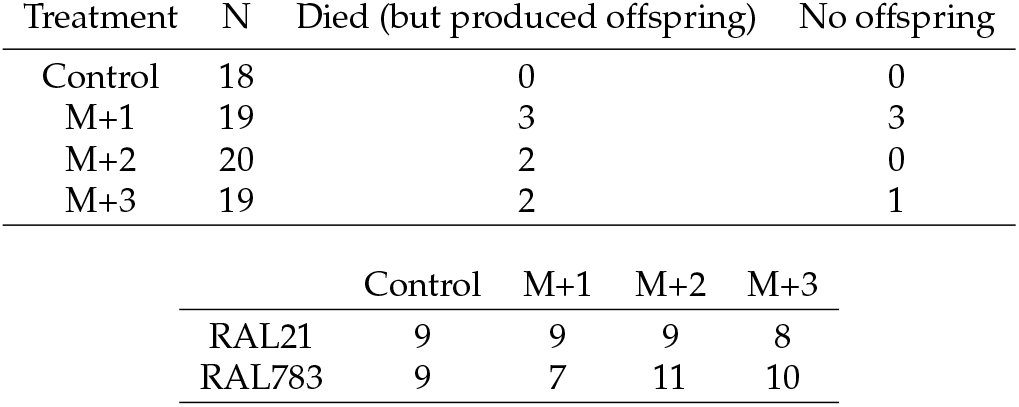

| Treatment | N | Died (but produced offspring) | No offspring |
| --- | --- | --- | --- |
| Control | 18 | 0 | 0 |
| M+1 | 19 | 3 | 3 |
| M+2 | 20 | 2 | 0 |
| M+3 | 19 | 2 | 1 |

|  | Control | M+1 | M+2 | M+3 |
| --- | --- | --- | --- | --- |
| RAL21 | 9 | 9 | 9 | 8 |
| RAL783 | 9 | 7 | 11 | 10 |

#### 5.2.3 Full model results – ANOVA

|  | Df | Deviance | Resid. Df | Resid. Dev | F | Pr(>F) |
| --- | --- | --- | --- | --- | --- | --- |
| \| NULL | \| NA | \| NA | \| 53 | \| 1575.115 | \| NA | \| NA |
| \| WOUND_TIMING | \| 2 | \| 34.7041 | \| 51 | \| 1540.410 | \| 0.7575 | \| 0.4748 |
| \| GENOTYPE | \| 1 | \| 222.1107 | \| 50 | \| 1318.300 | \| 9.6964 | \| 0.0032\|* |
| \| MATING_DURATION | \| 1 | \| 44.9535 | \| 49 | \| 1273.346 | \| 1.9625 | \| 0.1683 |
| \| FUNGUS | \| 1 | \| 35.5011 | \| 48 | \| 1237.845 | \| 1.5498 | \| 0.2198 |
| \| WOUND_TIMING:GENOTYPE | \| 2 | \| 1.1674 | \| 46 | \| 1236.678 | \| 0.0255 | \| 0.9749 |
| \| GENOTYPE:MATING_DURATION | \| 1 | \| 119.6440 | \| 45 | \| 1117.034 | \| 5.2232 | \| 0.0272\|* |
| \| GENOTYPE:FUNGUS | \| 1 | \| 19.7353 | \| 44 | \| 1097.299 | \| 0.8616 | \| 0.3584 |

TIME TO 50% ~ WOUND\_TIMING\*GENOTYPE + GENOTYPE\*MATING\_DURATION + GENOTYPE\*FUNGUS, family = "quasipoisson"
|  | Df | Deviance | Resid. Df | Resid. Dev | F | Pr(>F) |
| --- | --- | --- | --- | --- | --- | --- |
| \| NULL | \| NA | \| NA | \| 53 | \| 48.2737 | \| NA | \| NA |
| \| WOUND_TIMING | \| 2 | \| 2.3331 | \| 51 | \| 45.9406 | \| 1.2419 | \| 0.2988 |
| \| GENOTYPE | \| 1 | \| 0.4206 | \| 50 | \| 45.5200 | \| 0.4477 | \| 0.5069 |
| \| MATING_DURATION | \| 1 | \| 1.9807 | \| 49 | \| 43.5393 | \| 2.1085 | \| 0.1536 |
| \| FUNGUS | \| 1 | \| 0.5097 | \| 48 | \| 43.0296 | \| 0.5426 | \| 0.4653 |
| \| WOUND_TIMING:GENOTYPE | \| 2 | \| 2.4745 | \| 46 | \| 40.5551 | \| 1.3171 | \| 0.2783 |
| \| GENOTYPE:MATING_DURATION | \| 1 | \| 1.1605 | \| 45 | \| 39.3946 | \| 1.2354 | \| 0.2724 |
| \| GENOTYPE:FUNGUS | \| 1 | \| 2.7213 | \| 44 | \| 36.6733 | \| 2.8969 | \| 0.0958\|. |

TIME TO 95% ~ WOUND\_TIMING\*GENOTYPE + GENOTYPE\*MATING\_DURATION + GENOTYPE\*FUNGUS, family = "quasipoisson"
|  | Df | Deviance | Resid. Df | Resid. Dev | F | Pr(>F) |
| --- | --- | --- | --- | --- | --- | --- |
| \| NULL | \| NA | \| NA | \| 53 | \| 149.7527 | \| NA | \| NA |
| \| WOUND_TIMING | \| 2 | \| 1.6084 | \| 51 | \| 148.1442 | \| 0.3296 | \| 0.7209 |
| \| GENOTYPE | \| 1 | \| 9.7924 | \| 50 | \| 138.3518 | \| 4.0138 | \| 0.0513\|. |
| \| MATING_DURATION | \| 1 | \| 5.3737 | \| 49 | \| 132.9781 | \| 2.2026 | \| 0.1449 |
| \| FUNGUS | \| 1 | \| 3.1075 | \| 48 | \| 129.8706 | \| 1.2737 | \| 0.2652 |
| \| WOUND_TIMING:GENOTYPE | \| 2 | \| 4.7487 | \| 46 | \| 125.1219 | \| 0.9732 | \| 0.3858 |
| \| GENOTYPE:MATING_DURATION | \| 1 | \| 9.3920 | \| 45 | \| 115.7299 | \| 3.8496 | \| 0.0561\|. |
| \| GENOTYPE:FUNGUS | \| 1 | \| 0.0198 | \| 44 | \| 115.7101 | \| 0.0081 | \| 0.9286 |

OFFSPRING DIFFERENCE ~ (INJURY\_STATUS + DAY + GENOTYPE)^3
|  | Df | Deviance | Resid. Df | Resid. Dev | F | Pr(>F) |
| --- | --- | --- | --- | --- | --- | --- |
| \| NULL | \| NA | \| NA | \| 106 | \| 2914.897 | \| NA | \| NA |
| \| INJURY_STATUS | \| 1 | \| 266.5059 | \| 105 | \| 2648.391 | \| 11.2714 | \| 0.0011\|* |
| \| DAY | \| 2 | \| 207.9200 | \| 103 | \| 2440.471 | \| 4.3968 | \| 0.0149\|* |
| \| GENOTYPE | \| 1 | \| 14.5973 | \| 102 | \| 2425.874 | \| 0.6174 | \| 0.4340 |
| \| INJURY_STATUS:DAY | \| 2 | \| 45.1615 | \| 100 | \| 2380.713 | \| 0.9550 | \| 0.3885 |
| \| INJURY_STATUS:GENOTYPE | \| 1 | \| 18.6513 | \| 99 | \| 2362.061 | \| 0.7888 | \| 0.3767 |
| \| DAY:GENOTYPE | \| 2 | \| 17.9374 | \| 97 | \| 2344.124 | \| 0.3793 | \| 0.6854 |
| \| INJURY_STATUS:DAY:GENOTYPE | \| 2 | \| 97.9112 | \| 95 | \| 2246.213 | \| 2.0705 | \| 0.1318 |

#### 5.2.4 Full model results, all interactions – ANOVA

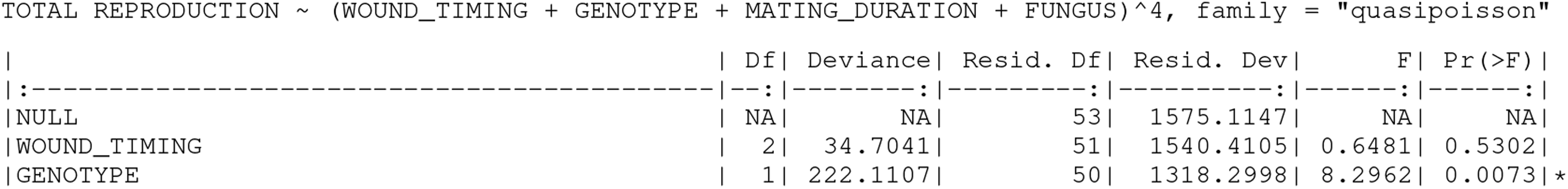
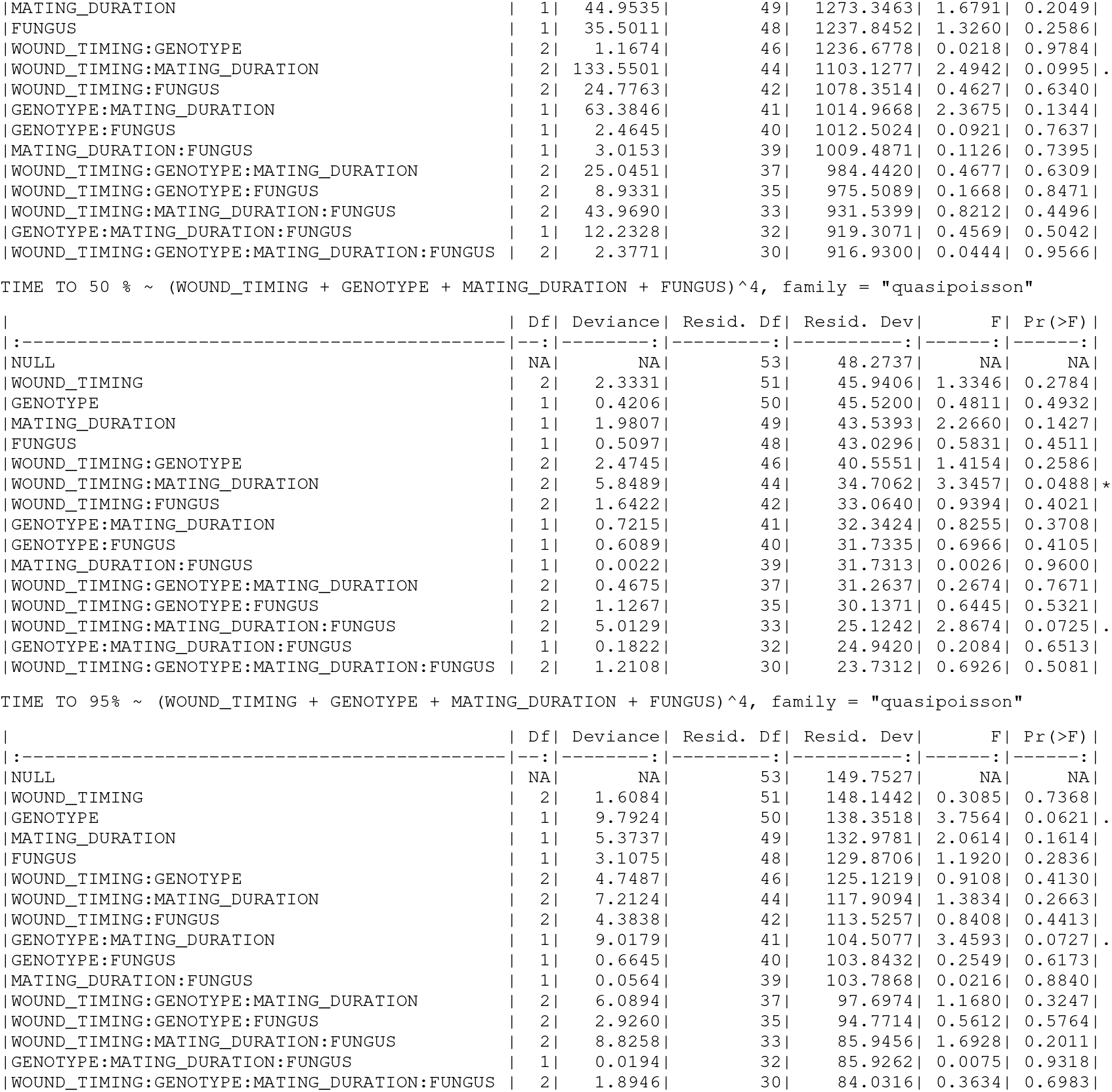

